# Deep Generative Design of Epitope-Specific Binding Proteins by Latent Conformation Optimization

**DOI:** 10.1101/2022.12.22.521698

**Authors:** Raphael R. Eguchi, Christian A. Choe, Udit Parekh, Irene S. Khalek, Michael D. Ward, Neha Vithani, Gregory R. Bowman, Joseph G. Jardine, Po-Ssu Huang

## Abstract

Designing *de novo* binding proteins against arbitrary epitopes using a single scaffold, as seen with natural antibodies, remains an unsolved challenge in protein design. Current design methods are unable to capture the structural dynamics of flexible loops nor search loop conformational space in a principled way. Here we present Sculptor, a deep generative design algorithm that creates epitope-specific protein binders. The Sculptor algorithm constitutes a joint search over the positions, interactions, and generated conformations of a fold, and crafts a backbone to complement a user-specified epitope. Sequences are designed onto generated backbones using a combination of a residue-wise interaction database, a convolutional sequence design module, and Rosetta. Instead of relying on static structures, we capture the local conformational landscape of a single fold using molecular dynamics, and demonstrate that a model trained on such dense conformational data can generate backbones tailor-fit to an epitope. We use Sculptor to design binders against a conserved epitope on venom toxins implicated in neuromuscular paralysis, and obtain a multi-toxin binder from a small naïve library – a promising step towards creating broadly neutralizing binders. This study constitutes a novel application of deep generative modeling for epitope-targeted design, leveraging conformational dynamics to achieve function.

## Introduction

Numerous essential biochemical processes and cell behaviors are regulated by protein-protein interactions (PPIs), and many studies have shown that engineered proteins targeting protein-protein-interfaces serve as effective therapeutics, powerful modulators of cell signaling [1, 2, 3, 4, 5, 6], and crucial components in recent CAR-T cell therapies [7, 8, 9]. Attesting to the need for binding proteins is the sheer scale of the biologics industry, which was valued at $300 billion in 2020, and expected to grow to $500 billion by 2026 [10]. Despite both the demand and utility of epitope-specific binders, engineering them remains a long-standing challenge, with most methods requiring screening of massive combinatorial libraries, requiring extensive effort to direct contacts towards the target epitope.

Computational protein design has enabled the creation of novel folds and a wide variety of de novo scaffolds [11, 12, 13, 14, 15, 16, 17, 18]. However, design of epitope-specific binders has remained difficult due to the need to create a foldable protein with both backbone and sequence that complement the epitope of interest. In recent work, denoising diffusion probabilistic models [19] have gained attention in protein design [20, 21, 22], and have been shown as being able to generate static helical binders against hotspots by using a variant of RoseTTAFold fıne-tuned as a denoising network [23]. A general method is RifDock, which Cao et. al. used to create protein-protein binders from a large collection of pre-built miniproteins. RifDock uses a hierarchical search scheme to dock rigid-body backbones into a rotamer interaction field [11] [24]. Polizzi et. al. reported a conceptually similar method called COMBs [25]. Similar to RifDock, the COMBs approach uses interaction units called “van der Mers” to identify valid backbone geometries from a set of pre-constructed helical bundles.

While interaction field methods are powerful, they depend on the creation of sufficiently large rigid-backbone libraries, and little is reported about how to maximize library compatibility against a wide variety of epitope shapes. While parametrically generated helical bundles can finely sample conformations, finding parameters that satisfy interaction-field constraints is non-trivial, and this approach does not generalize to non-helical structures. There is not yet a reliable way to design interfaces involving flexible loops such as those found in antibodies, and we speculate that dense sampling of the loop-conformation space during design is key to creating such interfaces. We hypothesize that a generative model capturing the fine-grained conformational space around a single fold would be able to tailor novel structures to fit design constraints, such as an interaction field, while incorporating structural dynamics into the design process.

In this study we combine interaction-field-guided design and deep generative modeling to perform flexible-backbone, epitope-specific binder design in a highly generalizable way. We build upon the generative design paradigm described in a prior study [26] creating an algorithm that “sculpts” a protein backbone to fit a target surface. Specifically, we use a protein VAE to capture a massive space of possible binder conformations sampled using molecular dynamics (MD) simulations. Then, given a target epitope, we dynamically assign interacting positions between both the target and binder, while optimizing the binder’s structure in the latent space to fit the interaction field. We design sequences using a combination of a neural-network-based design module [27] and Rosetta [28, 29, 30, 31, 32]. To experimentally validate our method, we challenge our algorithm to create a binder against a conserved motif on long-chain snake-venom toxins implicated in venom-induced skeletal muscle paralysis. We limit ourselves to a small library of ~5800 designs – far smaller than conventional yeast display libraries which are often ~ 10^7^ sequences or more – but are nonetheless able to generate a binder against the desired epitope and achieve binding across multiple venom toxins, a first step towards broad neutralization. Our algorithm, named “Sculptor,” constitutes one of the few experimentally-validated generative design methods, and is unique in that it combines a large amount of conformational data from molecular-dynamics simulations with deep generative modeling to create protein backbones with complementarity against user-specified epitopes.

## Overview of the Sculptor Algorithm

In order to design a binding protein that is complementary to the target interface at both the sequence and backbone structure level, there are four spaces that must be explored during design: (1) the space of rotational and translational degrees of freedom, (2) the space of backbone structure, (3) the space of interface assignments, and (4) the space of amino acid sequences. The core of our algorithm focuses on joint optimization over spaces (1) to (3), while (4) is searched using a combination of a learned sequence design module and Rosetta interface optimization after a backbone is created.

An overview of the Sculptor algorithm is shown in Figure 1. The input to Sculptor is a protein structure and a user-specified epitope. An interaction field is constructed around the epitope using an amino-acid specific interaction database consisting of pairwise interactions. Given a target field, the algorithm randomly initializes the latent vector of a coordinate-generating VAE that defines the backbone conformational search space[26]. Homogenous transformation parameters that dictate the position of the binder in 3D space, and an assignment of interacting residue pairs between the target and binder are also initialized. The core loop of Sculptor jointly optimizes all of these parameters to maximize interaction field fit using a combination of gradient descent, Metropolis-Hastings, and linear sum assignment (Figure 1, blue). Over the span of the trajectory, the initialized binder is molded against the target interface as it is fitted into the interaction field. We refer to this process as “Sculpting.”Once the protein backbone is sculpted, sequence design is done using a convolution-based sequence design module in combination with Rosetta. Residue choices from both the interaction field and the design module are passed to Rosetta which builds explicit side-chains and searches for jointly compatible sequences. In the final design step, Rosetta is allowed to freely redesign the sculpted interface to optimize interactions. A detailed description of the algorithm is provided in the Methods and Supplemental Methods section.

**Figure 1:**
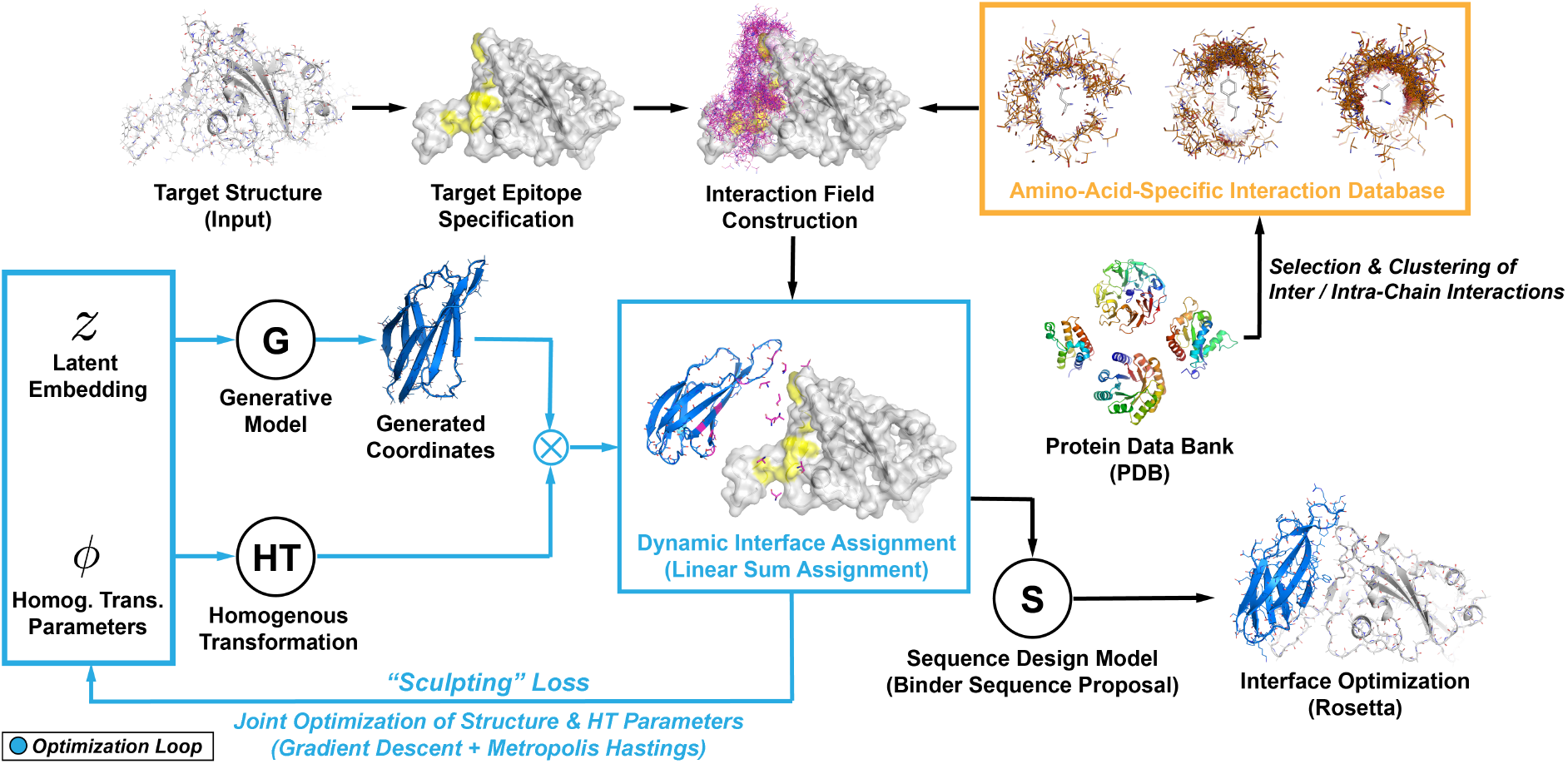
The Sculptor Algorithm. The input to sculptor is a target structure with a user-specified target epitope (yellow). An amino-acid-specific interaction field (magenta) is built around the target epitope using a database of clustered protein interactions harvested from PDB (orange). To perform interface design, a generated structure is docked into the interaction field while jointly optimizing the binder backbone conformation and position. During optimization the interacting residues on both the binder and target are dynamically reassigned via linear sum assignment to minimize fitting loss. We refer to this process, shown in blue, as “Sculpting.” Post-sculpting, the optimized structure is passed to a neural network-based sequence design module which provides homology-informed sequences that are combined with field-specified residues to propose candidate amino acids at each position. Final unrestricted interface optimization is performed using Rosetta.

## Components of the Sculptor Algorithm

### Deep Generative Model of Backbone Structure

The core component of Sculptor is a deep variational autoencoder trained similarly to the previously reported IgVAE [26], and provides a conformational search space for our design algorithm. Crucially, this model generates 3D coordinates that are fully differentiable with respect to the latent vector, allowing for computation of gradients and structural optimization via latent space search.

In this study we select the monobody scaffold as our design template. Monobodies are a family of binding proteins based on a modified fibronectin-3 scaffold [33, 34], and have many publicly available high-resolution structures in complex with a wide range of targets [1, 2, 3, 35]. Most reported monobodies are of similar length and have a conserved core, but have diverse loop sequences and structures. These features are well-suited to sequence design module and generative model training because most variance in the data is directly relevant to target specificity. To train the generative model, we selected 34 monobody PDBs of similar length and structurally padded these to be length 91 using Rosetta Remodel [32] as previously described [26]. In addition, we randomly generated 12 triple-mutants per structure, yielding a total of 442 seed structures. To create our training datasets, we simulated the seed structures using molecular dynamics (MD). Frames from the simulation were clustered and used to create a training set of 1.6M structures, yielding a dense conformational dataset biased towards native structures and augmented with physically realistic structures (see Methods).

In Figures 2A and 2B we analyze the performance of the generative model, showing backbone and torsional distributions of the training set and generated monobody structures. While slightly higher in variance, the generated structures are high in quality with the expected conformational and torsional propensities. These results are in agreement with prior characterizations of model performance [26], and suggest that the VAE adequately captures monobody conformational space.

**Figure 2:**
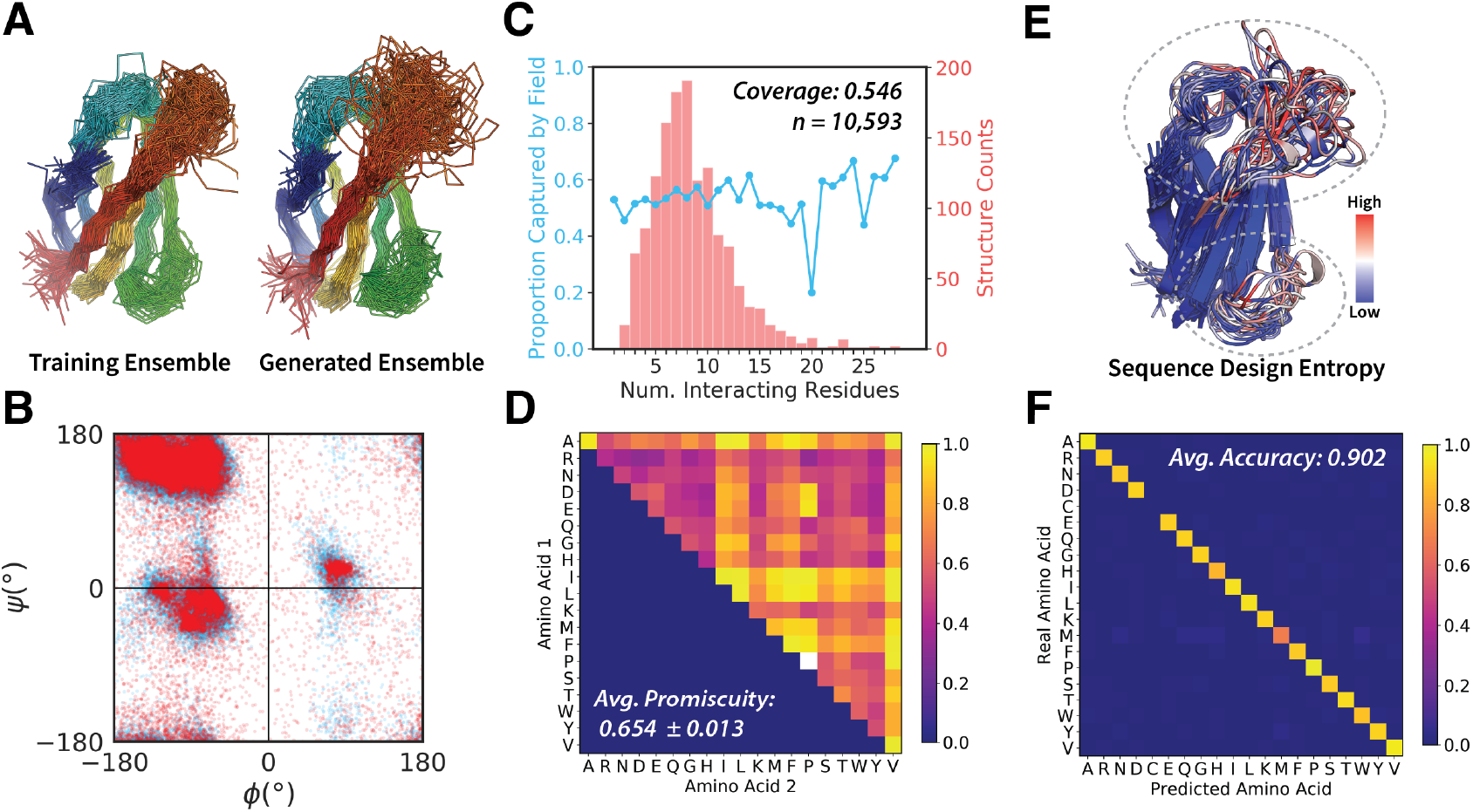
Performance of Algorithm Components. (A) Comparison of 100 randomly selected training set structures and 100 random VAE-generated structures. (B) Comparison of Ramachandran distributions of the training and generated structures. (C) Assessment of field coverage using the PPI4DOCK benchmark set. PPI4Dock contains 1,417 complexes with 10,593 amino acid interactions detected by Arpeggio. The proportion of residues captured by our field versus the number of interacting residues in a complex is plotted in blue with standard errors. The distribution of numbers of interacting residues in PPI4DOCK complexes is shown as a histogram in red. (D) Promiscuity matrix of the interaction field. Each matrix position corresponds to a pair of amino acid types, and the entry is the proportion of interactions for which there is a similar interaction within 0.5 Å dRMSD formed by other type-pairs. dRMSD is computed for backbone atoms (N, Ca, C, O, Cb) between the two interacting residues. (E) Position-wise entropy of sequence-design-module-predicted amino acid distributions for 30 generated backbones. Monobody binding loops are circled in gray. (F) Test set confusion matrix for the deep sequence design module. Each amino acid type is represented by 100 randomly selected examples, with the exception of cysteine, which are not present in native monobodies. Counts are normalized horizontally by the number of real examples in each class.

### Pairwise Residue Interaction Field

In Figure 2C and 2D, we show an analysis of the interaction database, which is used to construct an interaction field from a user-specified epitope. These interactions are harvested from the protein data bank (PDB) using Arpeggio [36], and clustered via hierarchical clustering. The database includes both inter-chain and intra-chain polar and hydrogen bonding interactions (See Methods). Figure 2C shows coverage of our field on the PPI4Dock dataset which contains 1,417 complexes and is widely used to benchmark protein-protein docking algorithms. We define the coverage threshold as having backbone atom (N, Ca, C, O, Cb) alignment within 0.5Å dRMSD [37]. The distribution of the number of interacting residues per complex is shown as a red histogram, and the percentage of interacting residues captured by our field in blue. Overall our field exhibits a coverage of 55%, and this value remains relatively constant across different interface sizes. Importantly, this field coverage value does not indicate field quality, but rather that our design space is comprised of the 55% most enriched interactions in PDB.

Figure 2D depicts the “promiscuity matrix” of our interaction field. Each entry in this matrix indicates the proportion of interactions for a given amino acid pair that are geometrically similar to any other amino acid pair with different identities. We refer to this as the promiscuity value. For example, for the pair (V,V) the entry value is close to 1.0. This indicates that nearly all (V,V) interactions have some non-(V,V) pair in the database that is very similar in relative positioning, which we define as having backbone atom alignment within 0.5Å dRMSD. Notably, many residue pairs had a high promiscuity, with an overall average of 65.4%. This suggests that many residue-residue interactions adopt highly similar 3D positions independent of residue identity. This observation motivates the final step in our pipeline after sequence design, where Rosetta is allowed to freely redesign the interaction interface, since many field-specified pairs may have alternate amino acid identities that form favorable interactions.

### 3D-Convolutional Sequence Design Module

Sculptor’s sequence design scheme is built with the intent to restrict the sequence search space by biasing designs towards known monobody sequences while also incorporating interaction-field-specified amino acid identities. Post backbone-sculpting, we use a 3D-convolution-based neural network to perform sequence design. Similar to our previously described method [27], the design module is trained as a classifier that takes a voxelized residue environment as input, and outputs a probability distribution over amino acid identities. Differently from the published method, Sculptor’s design module uses only backbone atoms to make side-chain-independent predictions at each position. This allows the model to capture backbone-position-dependent homology specific to the monobody fold, rather than being a general protein design tool. Importantly, the design module is used only to specify the sequence of the binder and does not “see” any features on the target epitope – these are instead handled by the interaction field. The outputs of the design module are used to create a preliminary list of candidate amino acids at each position by selecting residue types above a fixed probability threshold. These candidates are combined with the fıeld-specified interface residues before being passed to Rosetta for joint design and repacking at all positions. By structuring the sequence design scheme this way, field-specified positions not used in the final interface design are biased towards a native-like distribution.

In Figure 2E we visualize the residue-wise Shannon entropies from the sequence design module for 30 randomly generated backbones. The entropy of the amino acid distributions are lower along core beta-strands and reflect the conserved sequence identities found in existing monobodies. Entropies are noticeably higher in the two side-loops, which are used to bind a variety of targets and have the most variable sequences. This result suggests that the design module correctly captures uncertainty in monobody sequence space and its backbone-positional dependence. Design module accuracy and a confusion matrix are shown in Figure 2F. The module achieves an average accuracy of 90.2% over a balanced test set of 100 examples per amino acid type. Confusion between different residue types is low (< 0.2) overall. Further details regarding sequence design module training are provided in the Methods section.

## Recovery and Redesign of a Native Complex

To assess the functionality of Sculptor we tested to see whether it could recover the native structure of a monobody in an existing complex. As a test case we selected PDB:2OCF(A), which is the structure of a monobody bound to the human estrogen receptor ligand-binding domain [35]. This complex was selected because it uses a helical binding loop, which is rare among known monobodies, making it both underrepresented in our training dataset and more difficult to recover via latent space search.

To test recovery of the 2OCF monobody, we specified the native epitope as the target interface and sampled 5,864 sculpting trajectories each initialized with a random backbone conformation and random position centered around the target protein. The trajectory that recovered the native complex most closely is shown in Figures 3A and 3B and achieves a native RMSD of 1.2Å. Inspection of the trajectory suggests the algorithm traverses a large span of conformational space before it is molded against the target epitope and shaped into a helical structure (Figure 3A). After sequence design and unconstrained refinement with Rosetta, native RMSD improved to 0.97Å. Notably, pre- and post-refınement outputs exhibited very little change in backbone structure (< 1Å RMSD), suggesting the complex recovered by Sculptor likely corresponds to a local energy minimum over the conformational and positional space explored by the algorithm. Post-design, the sequence identity of the recovered binder was 52.7% relative to native. While sequence recovery was somewhat low, the ddG of recovered and native sequences were both approximately −50 Rosetta Energy Units (REU), indicating that the alternative sequence is favored similarly to native.

**Figure 3:**
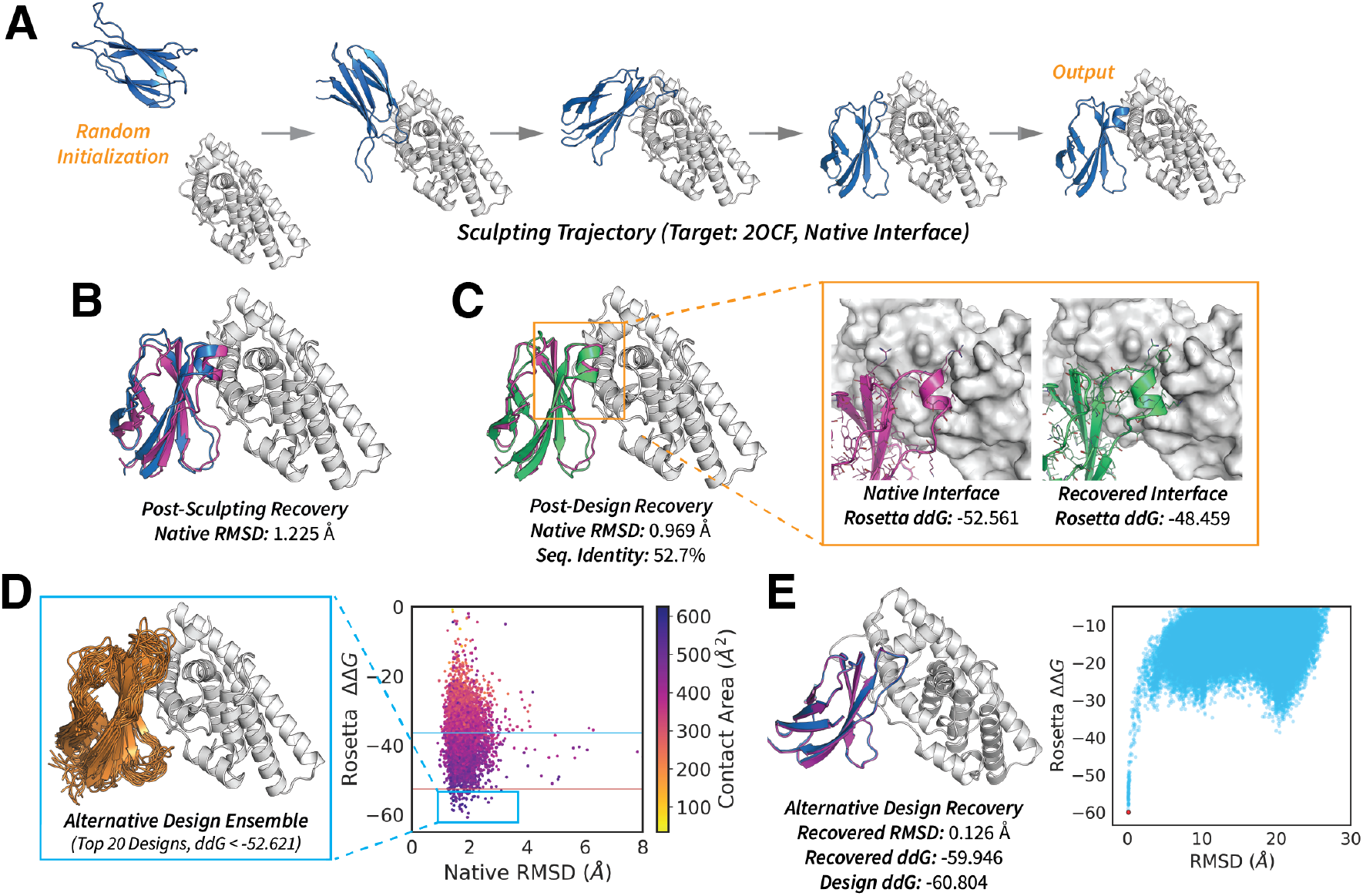
Recovery and Redesign of a Native Complex. (A) Sculpting trajectory against 2OCF(A) given the native interface assignment. The generated molecule is shown in blue, the target is shown in white. (B) Alignment of the post-sculpting structure, shown in blue, and the native complex, shown in magenta. (C) Alignment of the post-sequence-design structure, shown in green, and the native complex, shown in magenta. The insert depicts the full-atom rendering of the helical loop from each structure interacting with the target surface. (D) Plot of ddG’s of designs generated with the native 2OCF interface assignment. The red line is drawn at the ddG of the native complex (−52.621 REU) and the blue line is drawn at the average ddG of all complexes in the PPI4DOCK dataset (−36.437 REU, n=1,417). The insert depicts the ensemble of top 20 designs with ddG values better than the native complex. (E) Docking recovery of the top alternative design from the ensemble shown in panel D. The designed structure is shown in magenta and the lowest ddG structure from the docking trajectory is shown in blue. The full trajectory (n=100,000 decoys) is shown in the plot on the right. The lowest ddG structure is shown as a red point.

Next, for the same epitope we asked whether our algorithm could provide novel binder designs that differed from the native binder. In Figure 3D we show an ensemble of the top 20 designs ranked by ddG and show a plot of ddG versus native RMSD. Among the same set of 5,864 trajectories, we found that our algorithm generated 2,977 interfaces with ddG’s better than the average ddG of PPI4DOCK (Figure 3D, blue line), among which 68 interfaces achieved ddG’s better than the native 2OCF interface (Figure 3D, red line). To further test epitope specificity we performed blind global docking of the best scoring complex with RosettaDock [38, 39], with results shown in Figure 3E. The designed complex was recovered to within 1Å RMSD, and the docking trajectory formed a robust funnel converging to a global energy minimum over 100,000 decoys. Overall these results suggest our algorithm is able to recover the backbone structure of a native complex via the sculpting process, even when the loop structure is poorly represented. Further, Sculptor is able to provide designs that differ from known native binders, but are comparable in interface quality under Rosetta evaluation.

## Computational Validation of Novel Epitope-Specific Designs

To assess algorithm performance against epitopes for which monobody binders do not exist, we selected 3 targets: SHV-1 beta-lactamase PDB:3C4P(A), alpha-elapitoxin PDB:4LFT(B), and SARS-CoV2-RBD PDB:6VW1(E). For each of these targets we chose epitopes identified in literature as being biologically relevant (Figure 4, yellow). The beta-lactamase epitope corresponds to a known inhibitory site [40], the toxin epitope corresponds to a motif on long-chain snake venom toxins needed for nicotinic acetylcholine receptor (nAChR) binding [41], and the RBD epitope corresponds to the ACE2 binding site [42]. We used Sculptor to create 3,000 designs against each epitope, and selected structures with the best ddG’s from each set. In each trajectory the monobody structure was initialized at a random position without any contacts to the target epitope before undergoing sculpting. In Figure 4 we used blind global Rosetta docking was used to computationally verify design specificity. We found that all three examples closely recovered the designed interface with docking trajectories forming robust funnels, and designed binding modes constituting global energy minima over 100,000 decoys (Figure 4 plots, red dots). Similar to the 2OCF designs, post-sculpting backbones moved very little after unconstrained design (< 1Å RMSD), suggesting again that post-sculpting designs appear adopt a local energy minima under Rosetta (Figure 4, Trajectory vs Designed).

**Figure 4:**
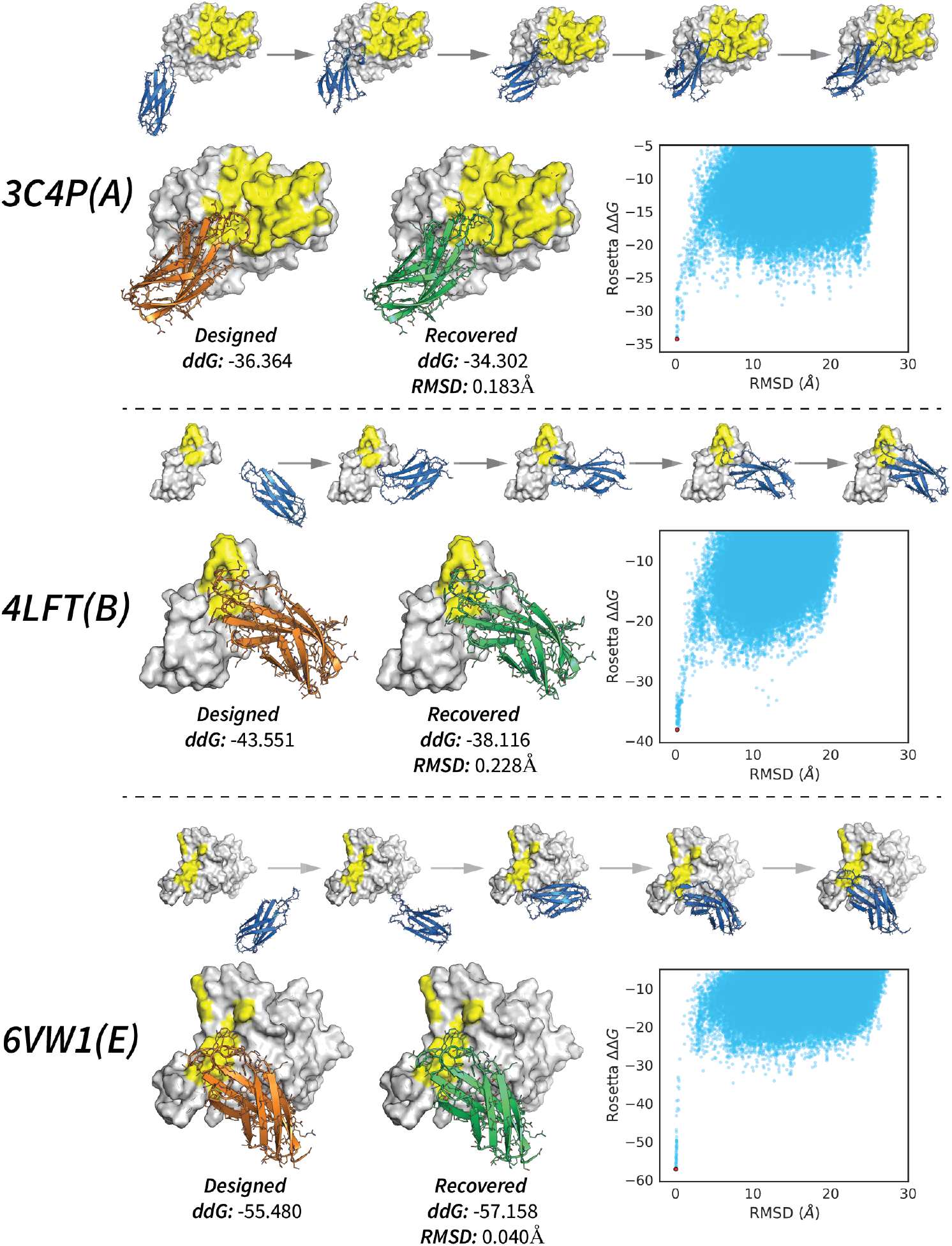
Sculpting Trajectories and Blind Recovery of Designs. The top sequence of structures depicts frames from the sculpting trajectory for each target starting from a random initialization. Targets are shown in white with epitopes colored yellow. The orange structures correspond to the full-atom designs. The green structures are the structures recovered from blind global docking using RosettaDock. The docking trajectories are shown in right-side plots, with each run containing 100,000 decoys and minimum energy decoy shown as a red point.

While the sculpting process constitutes a search over possible interaction modes within the designated epitope, we note that potential binders are allowed to form any number of interactions and are not required to mask the entire epitope. This flexibility allows the algorithm to adapt when the user specifies a very difficult-to-reach epitope, for example, by using only a small portion of a binding loop for interaction. This effect is observable in differences in epitope coverage between the three examples, where Sculptor places a loop in a groove on the surface of 3C4P(A), while leaving the right side more exposed. In contrast, the designs against 4LFT(B) and 6VW1(E) cover a larger proportion of the specified surface. This behavior can be controlled by the user by simply adjusting the size of the specified epitope. Overall our data suggest that Sculptor is able to generate realistic binders against a variety of epitopes that are favorable under computational benchmarks. In addition, the interface-search functionality of the algorithm is robust, as it is able to recover promising binding modes from user-specified epitopes even when the specified area is large.

## Design and Experimental Characterization of a Multi-Toxin Binder

While Sculptor exhibits promising behavior under computational benchmarks, we sought to test our algorithm against a biologically significant and challenging epitope. We selected a conserved epitope on long-chain three-finger neurotoxins in snake venoms, implicated in their ability induce skeletal muscle paralysis via postsynaptic nAChR’s [41]. We sought to use Sculptor to create a pan-neutralizing binder that could be used to recognize this epitope across multiple venom toxins.

As target sequences we selected five toxin variants which contained the conserved epitope, labelling them T1 through T5 (Figure 5A). Crystal structures for T2, T3, and T4 were available on PDB and were used as inputs to sculptor. The structural overlay of these targets is shown in Figure 5B, with structures shown in white and the conserved motif shown in red. For each of the 3 targets, we sculpted ~8000 backbones, and designed sequences on backbones that fitted four or more field-interactions. The top 2000 binders for each target were selected using a combined ranking of Rosetta ddG, Sc [43], and interface-buried-SASA. This yielded a total library size of 5,923 designs against the common epitope. The full library was transformed into yeast and screened against all five of the targets. We were able to select and enrich for one binder designed against T3, and found that it bound to 3 of the 5 targets, specifically T3, T4 and T5, but did not bind to T1 or T2 (Figure 5B). The structure of the designed binder is aligned against the conserved epitope in Figure 5A, and the designed complex is shown in Figure 5D.

**Figure 5:**
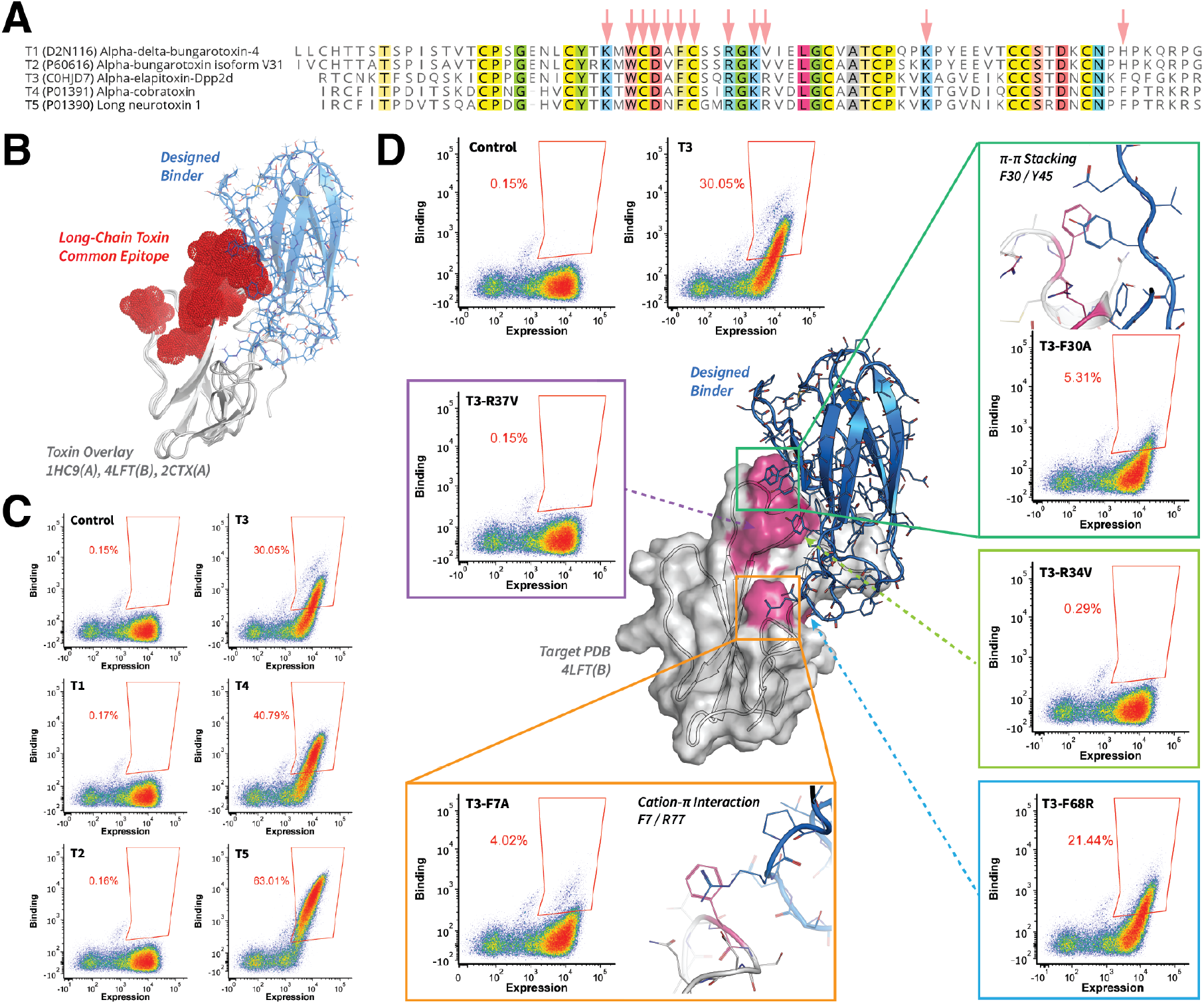
Epitope Mapping of a Sculptor-designed Venom Toxin Binder. (A) Sequence alignments of the five venom-toxin targets with gene accession numbers shown in parentheses. The target epitope is indicated by red arrows. (B) An overlay of the structures of three long-chain toxins with the common residues used for targeting shown in red. The overlaid PDBs correspond to T2, T3 and T4 respectively. (C) The binding profiles of the designed binder to the five targets. (D) The sculptor-designed 3D model of the enriched binder is shown in the center. The binder is shown in blue and target structure show in white. Each scatter plot shows a yeast surface display experiment analyzed with fluorescence assisted cell sorting (FACS). Display signal is plotted on the x-axis and binding signal on the y-axis. FACS gates are shown in red and annotated with binding population percentage. The two plots in the upper left correspond to binding profiles of the negative control and design target, T3. All other plots show binding profiles of mutants that attenuated binding relative to the native target. Attenuating mutants are mapped onto the target surface and colored magenta.

To perform epitope mapping, we introduced mutations the designed interface and found several that affected binding (Figure 5D). Two mutations, F7A and F30A reduced binding and two other mutations, R34V and R37V, abolished binding almost completely. These data suggest that our binder contacts the region comprised of F7, F30, R34, and R37, which is colored magenta on the designed complex in Figure 5C. The result shows a significant overlap between these mutations and the intended epitope, providing evidence in agreement with the design model.

To further confirm our binder’s epitope and understand cross-toxin specificity, we sought to identify the set of features in T3 required to rescue binding in T2. From sequence alignments, we observed that two residues, R37, and F66 were conserved across T3/4/5, but not in T1/2. While experiments in Figure 5D suggested that R37 is essential to T3 binding, we found that the equivalent mutation in T2 alone was not sufficient to rescue binding (Figure S1, T2 V37R). However, combining the V37R mutation with H66F allowed for recovery of binding (Figure S1, T2 V37R + H66F). From the crystal structure data, residue F66 residue appears to play a crucial role in determining the trajectory of the C-terminal tail (Figure S1B), and the rescue-effect suggests that the binder is making contact with F66 and likely other regions of tail as well. The importance of the C-terminal tail is further supported by the loss in binding when replacing the tail of T3 with that of T2 (Figure S1, T3 w/T2-CTerm). Interestingly, we discovered that V37R and H66F and were two of only three residues in that differed in the user-specified epitope between T3/4/5 and T1/2 (Figure 5A, red arrows), further suggesting that this feature is likely the main determinant of binder specificity in the native targets. We were also able to rescue T2 binding without the H66F mutation by replacing the first loop of T2 with that of T3, in combination with the V37R mutation (Figure S1, T2 w/T3-Loop1).

Overall our data suggest that the binder’s epitope agrees closely with with our design model and lies within the user-specified epitope. The T3 mutation data confirms that mutations at the interface result in loss of binding, and the T2 rescue data support this, suggesting that specificity to T3/5/6 is likely due to the combination of the R37 and F66 residues. The rescue experiments further suggest the binder contacts Loop 1, again in agreement with the T3 and the design model. Details of the yeast display experiments are provided in the Methods section. While it is likely necessary to affinity mature the Sculptor-designed binder for practical use, we note that the total library size used for these experiments is on the order of ~ 10^3^ and considerably smaller than that required by conventional yeast display libraries. Sculptor thus provides a powerful way to build small naïve libraries for obtaining initial binders for evolution – the main bottleneck in most protein engineering tasks – and a significant step towards broadly neutralizing binder design.

## Discussion

In this study we have presented a unique approach to epitope-specific binder design by leveraging dense conformational information collected through MD simulations. In contrast to other design methods, we focus on exploring the conformational space of a single fold, using its intrinsic flexibility to achieve function in a fashion similar to natural proteins. By posing epitope-specific design as a constrained optimization problem in the latent space of a generative model, we have created an algorithm that jointly docks and designs a molecule while incorporating loop conformational dynamics into the design process, tailor-fitting a protein structure to the interaction field constituting the space of favorable interactions. The Sculptor algorithm consolidates multiple ideas from state-of-the-art structure-based design methods into a coherent pipeline, drawing from our own work with the IgVAE [26] and convolutional sequence design [27] while also building on ideas from RifDock [11, 24], and COMBs [25]. Sculptor ultimately provides a new way to combine interaction field methods with deep generative models via dynamic interface assignment and latent structural optimization. From a practical standpoint, Sculptor is both modular and general since the generative model can be trained on any fold, allowing for broad choice of scaffold beyond monobodies, such as antibodies, nanobodies, ScFv’s, DARPins, lectins, and more. While the generative model in the Scultor algorithm is a VAE, our scheme is also compatible with other backbone generators, such as GANs and diffusion models.

The current iteration of Sculptor is designed for targeting arbitrary protein epitopes, but its core framework is highly flexible, and can be easily adapted for other molecule types such as nucleic acids and small molecules. In addition, Sculptor demonstrates a concrete application of the generative design paradigm, which allows us to solve arbitrary design tasks as a constrained optimization problem under a generative prior, so long as we can formulate a suitable objective function. Our work also reveals an important application of molecular dynamics in the age of deep learning, demonstrating that the conformational landscapes provided by simulation can be used to directly inform protein design, especially in tasks that require high conformational precision.

## Methods

### Algorithm

A detailed description of the Sculptor optimization loop and loss functions are provided in the Supplemental Methods. A movie of the Sculptor trajectory from Figure 3 is provided at tinyurl.com/sculptormb.

### Data Selection and Processing

Monobody structures that were similar in length (approx 91 residues) were manually chosen from PDB, yielding the following list of 34 structures: 6b2bC, 710gH, 5dc0A, 4jegB, 5e95B, 5g15B, 3csbA, 5dc4B, 6tlcD, 3csgA, 7jw7B, 6bqoC, 2obgA, 6o02B, 5kbnD, 6b2aD, 6apxB, 5a43D, 5ecjE, 7l0fB, 3qhtD, 5komD, 5dc9B, 6bynM, 5mtmB, 2ocfD, 3uyoD, 5n7eA, 5v7pD, 5mtjB, 4je4B, 7jxuD, 5mtnB, 6bx5D. Next, each template was structurally padded or truncated to 91 residues as described previously[26]. We randomly generated 12 triple mutants per structure using PyRosetta, with each mutation localized to the loop regions of the monobodies. Including the native PDBs, this yielded a total of 442 seed structures for simulation. The explicit list of mutants and their structures are available upon request.

### Data Generation using Molecular Dynamics

To generate training data we used molecular dynamics simulations. All simulations were performed using the AMBER03 force field for the proteins [44], in a dodecahedron solvated box using explicit TIP3P solvent model [45]. The solvated structures were energy minimized with a steepest descent algorithm using Gromacs 2020 [46], until the maximum force was below 100 kJ mol^-1^ nm^-1^. A step size of 0.01 nm and a cut-off distance of 1.2 nm for the neighbor list, Coulomb interactions and van der Waals interactions were used during the energy minimization. Following energy minimization, structures were equilibrated or 1 ns, and all bonds were constrained with the LINCS algorithm [47]. Virtual sites were used to enable an integration time step of 4 fs. A cut-off distance of 1.1 nm was used for the neighbor list, and a cut-off distance of 0.9 nm was used for Coulomb and van der Waals interactions. The Verlet cut-off scheme was used for the neighbor list. For the long-range electrostatic interactions, the particle mesh Ewald method was used with a Fourier spacing of 0.12 nm. Throughout the simulations, stochastic velocity rescaling (v-rescale) thermostat [48] was used to maintain the temperature of the system at 300 K.

Starting from the equilibrated structure for each of 442 monobodies, unbiased MD simulations were performed on the Folding@home distributed computing project [49]. Ten parallel simulations, each starting with randomly generated initial atomic velocities, were initiated from the equilibrated structures. An integration time step of 4 fs was used during these Folding@home simulations. The temperature of the system was maintained at 300 K using V-rescale thermostat and the pressure was controlled at 1 bar using Parrinello-Rahman barostat. Atomic coordinates of the protein were saved at every 20 ps time step in the Folding@home output trajectories, and an aggregate simulation time of more than 300 ns was collected for each monobody structure (164 us aggregate simulation). Folding@home output trajectories were further subsampled to retain every 5th frame in the trajectory. The resulting ~1.6M structures were used as training examples for the VAE. To create a dataset for the sequence design module, we randomly selected 40,000 of the clustered frames from the MD simulations and extracted the 3.64M residue environments corresponding to each residue in each frame. For the test set of the sequence design module we randomly selected 100 residue environments per amino acid type to ensure class balancing.

### Model Architectures and Training

The VAE architecture is reported in a prior study[26] with input and output dimensions adjusted to match the size of the structurally padded monobodies. In addition to the backbone atoms, the current model also includes Cb atoms, which are artificially placed for residues lacking them. Methods and scripts for structural padding are also provided in the IgVAE study [26].The architecture of the sequence design module is also similar to that previously reported[27], however the version implemented in this study only processes backbone atom coordinates (N, Ca, C, O), and Cb. This version of the model also does not construct side-chains, and performs classification at each position, independent of amino acid identities at other positions. The residue environments used as inputs are 8Å voxelized cubes centered at the residue-of-interest and with a voxel size of 0.5Å. All models were trained on RTX 2080Ti’s using the PyTorch deep learning framework [50].

### Sequence Design

Sequence design is performed in two phases. In the first, a list of interacting residues is created using nearest alignments to the interaction field and merged with the list of design-module-proposed amino acid identities. Any amino acids with greater than ~ 5% probability under the classifier are included as candidates. The combined list is passed to Rosetta for joint sequence design at all positions. During this phase a cartesian relax is performed during design, allowing for optimization of bond lengths and angles. In the second phase, Rosetta is used to optimize the binding interface by designing without restrictions to amino acid identity, and the designed structure undergoes an unconstrained relax. The first phase uses the PyRosetta [30] interface, while the second uses a RosettaScripts [29] interface.

### Interaction Field Construction

Protein structures were downloaded from RCSB Protein Data Bank with 40% sequence homology, X-ray diffraction resolution ≤ 2.0 Å, and *R_obs_* ≤ 0.2. This resulted in a list of 14,680 structures. Each structure was analyzed using Arpeggio [36] to identify all residues involved in polar interactions with interaction distances ≤ 5 Å. Interacting residues on the same PDB chain are required to be more than 6 residues apart. The interacting residues are grouped by amino acid identity resulting in 210 groups (e.g. Ala-Ala, Ala-Arg,…). For each group an all-vs-all distance matrix is computed using RMSD. The backbone atoms (N, C, Ca, O) and Cb atom was used for the RMSD calculations. For glycine a virtual Cb atom was generated. In groups where both interacting amino acids were the same identity (e.g. Ala-Ala, Arg-Arg,…) RMSD was computed twice by flipping residue order, with the minimum of the two being used as the final value. Clusters were created using hierarchical clustering on the distance matrix with an distance threshold of 0.3Å dRMSD and ward linkage to minimize variance within each cluster. For each cluster, the representative pair was determined by the pair that minimizes the sum of the distances to all other pairs in the same cluster.

### Yeast Display Experiments

#### Recombinant Production of Three Finger Toxins

Long chain alpha-neurotoxin three finger toxin (3FTX) variant sequences (Table S2) from various snake species were selected from NCBI databases and cloned into a mammalian expression vector containing a C-terminal Avi™ tag followed by a WELQut™ cleavage tag and a rabbit Fc tag by Genscript. Plasmid DNA was prepped and co-transfected with a plasmid expressing the BirA enzyme for in vivo biotinylation into Expi293 cells using FectoPRO^®^ (Polyplus Transfection). Cells were grown for 5 days shaking at 240 RPM, 37°C with 8% CO2 in Expi293™ expression medium (Thermo Fisher). Approximately 24 hours post-transfection, cells were stimulated with 3 mM valproic acid, 1.8 mg/ml D-glucose, and 20 nM d-biotin. Supernatant was harvested from cell cultures and incubated overnight rotating at 4°C with rProtein A Sepharose affinity resin (Cytiva). The affinity resin was washed once with 500 mM NaCl, 1X PBS and twice with 1X PBS before transferring to 20 mM Tris, 150mM NaCl, pH 7.4. WELQut protease (Thermo Fisher) was added to the resin and incubated with rotation for 3 hours at 30°C. Supernatant was harvested from the resin and incubated with HisPur™ Ni-NTA resin (Thermo Fisher) for 30 minutes at room temperature to remove the His-tagged protease, followed by a second incubation with Protein A resin for 30 minutes to remove any residual Fc tag. The final supernatant was concentrated to > 1 mg/ml using a 3 kDa centrifugal filter unit (Millipore Sigma) and frozen in aliquots stored at −80°C. Biotinylation of the purified 3FTX was confirmed using the Pierce™ biotin quantitation kit (Thermo Fisher). Tetramers of streptavidin conjugated to biotinylated 3FTX were formed via 60 min incubation of 4:1 3FTX:streptavidin-APC (Thermo Fisher) at room temperature prior to use.

#### Yeast Library Production and Transformation

We used Sculptor to generate 12,000 monobody designs against the common epitope for each of the target structures, PDB:1HC9(A), PDB:4LFT(B), and PDB:2CTX(A). The top 3,000 designs for each target were selected based on a combined ranking over Rosetta ddG, Sc, and buried SASA. Selected sequences were codon optimized for expressing in Saccharomyces cerevisiae and synthesized by Twist Biosciences. The pooled monobody libraries were PCR amplified and extended with long oligos to include 80bp of upstream homology and 55 bp of downstream homology with the pYDSI2u surface display vector. The amplified monobody encoding DNA and linearized pYDSI2u vector were transformed into yeast for homologous recombination.

#### Fluorescence-Activated Cell Sorting

In vitro engineering of antibodies or nanobodies can lead to constructs that are polyspecific[51], so both positive and negative selections were employed to enhance the specificity of the synthetic constructs. Yeast cells were alternatively selected as three-finger toxin (3FTX) binders (affinity sorts), or as poly-specificity reagent (PSR) non-binders (negative sorts). During affinity sorting, 1-5 × 10^7^ induced yeast cells were incubated for 60min rotating at 4°C with streptavidin-APC tetramerized 3FTX in PBSA (PBS containing 1% BSA) supplemented with 1 ug/mL anti-V5-AF405 to check the yeast display. During PSR sorting, induced yeast cells were incubated for 60 min rotating at 4°C with biotinylated HEK-cell soluble membrane protein extracts [52] in PBSA, washed with PBSA, then coupled to 1 ug/mL of both fluorophores (streptavidin-APC and anti-V5-AF405) for 20 min. Yeast cells were then washed once and resuspended in PBSA for sorting on a FACS Melody (BD Biosciences). Selected yeast cells were sorted into SD-Ura medium, grown shaking overnight at 30°C and induced for consecutive rounds of selection. Induction medium contained 20 g/L galactose, 1 g/L glucose, 6.7 g/L yeast nitrogen base, 5 g/L bacto-casamino acids, 38 mM disodium phosphate, 72 mM monosodium phosphate, and 419 uM L-tryptophan. For the 1st, 2nd and 3rd affinity sorts, 500, 500, and 20 nM of streptavidin-3FTX tetramer was used, respectively. For the single PSR sort (between the 2nd and 3rd affinity sorts) 20 ug/mL of biotinylated CHO-cell soluble membrane protein extract was used.

#### Monobody Sequencing and Analysis

Serial dilutions of the final affinity sort were plated on SD-Ura agar. After 3 days at 30°C, 30 single yeast colonies were combined and inoculated into 2 mL SD-Ura medium, and grown overnight shaking at 30°C. The DNA from the yeast cells was miniprepped (Qiagen) in the presence of zymolyase (Zymo Research) and transformed into DH10B competent E. coli (New England BioLabs). 30 single colonies were picked and analyzed via Sanger sequencing (Azenta Life Sciences). The same yeast culture was also analyzed via flow cytometry for binding to various 3FTX variants and mutants (Tables S2, S3) using 100 nM of streptavidin-APC tetramerized 3FTX (Figures 5, S1).

## Author Contributions

R.R.E., C.A.C and P.-S. H. conceived of the research. The codebase was written by R.R.E. and C.A.C., M.D.W. and N.V. ran molecular dynamics simulations and performed analysis and clustering of the trajectories. I.S.K. performed the yeast display experiments and identified the venom targets. U.P. performed experimental optimization of monobodies. The manuscript was written by R.R.E. and P.-S. H. with contributions from all authors. J.G.J., G.R.B., and P.-S.H. supervised the research.

## Acknowledgements

This project was supported by startup funds from the Stanford Schools of Engineering and Medicine, the Stanford ChEM-H Chemistry/Biology Interface Predoctoral Training Program, and the National Institute of General Medical Sciences of the National Institutes of Health under Award Numbers T32GM120007 and 1R01GM14789301. Additionally, this project was supported by the U.S. Department of Energy, Office of Science, Office of Advanced Scientific Computing Research, Scientific Discovery through Advanced Computing (SciDAC) program. DNA libraries were produced with support from Twist Bioscience. This work is also supported by funding from the Merck Research Laboratories SEEDS program.

---

**Figure S1:**
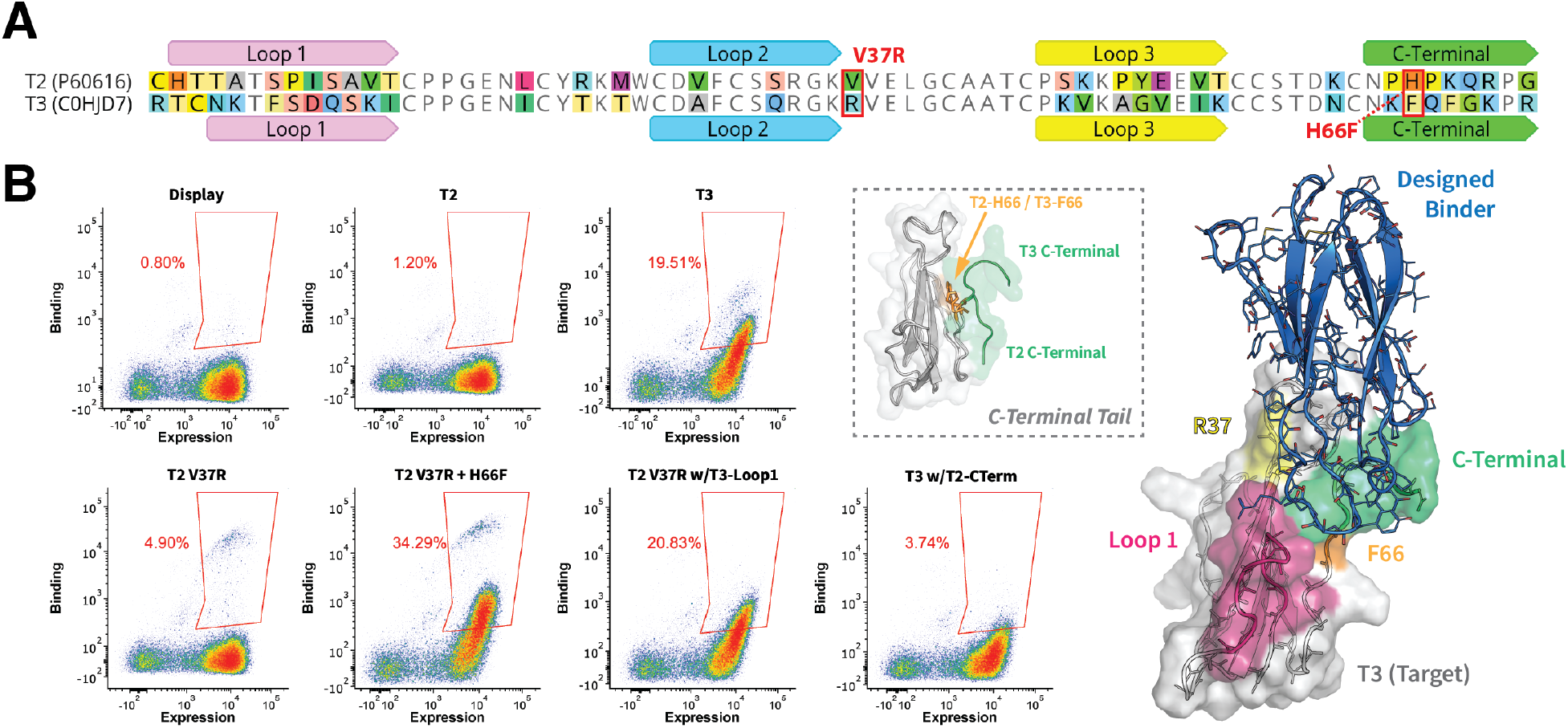
Rescue of T2 Binding using Designed Epitope Features. **(A)** Sequence alignments of T2 and T3 venom-toxin targets. Note the first two residues of T2 are truncated to denote equivalent positions more easily. Differing residues are highlighted and colored by type. **(B)** Each scatter plot shows a yeast surface display experiment analyzed with fluorescence assisted cell sorting (FACS). FACS gates are shown in red and annotated with the binding population percentage. The three plots in the top row correspond to a negative control, T2, and T3. Display signal is plotted on the x-axis and binding signal on the y-axis. The sculptor-designed 3D model of the enriched binder is shown on the right. The binder is shown in blue and target structure shown in white. Regions of interest are highlighted different colors. The dotted insert depicts a comparison of the difference in C-terminal tail structure between T2 and T3.

**Table S2:**
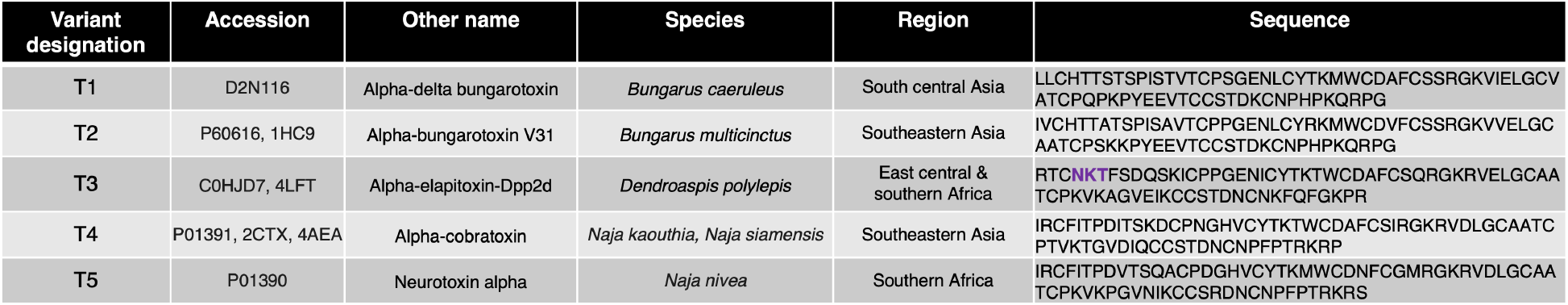
Three-Finger Toxin Sequences.

**Table S3:**
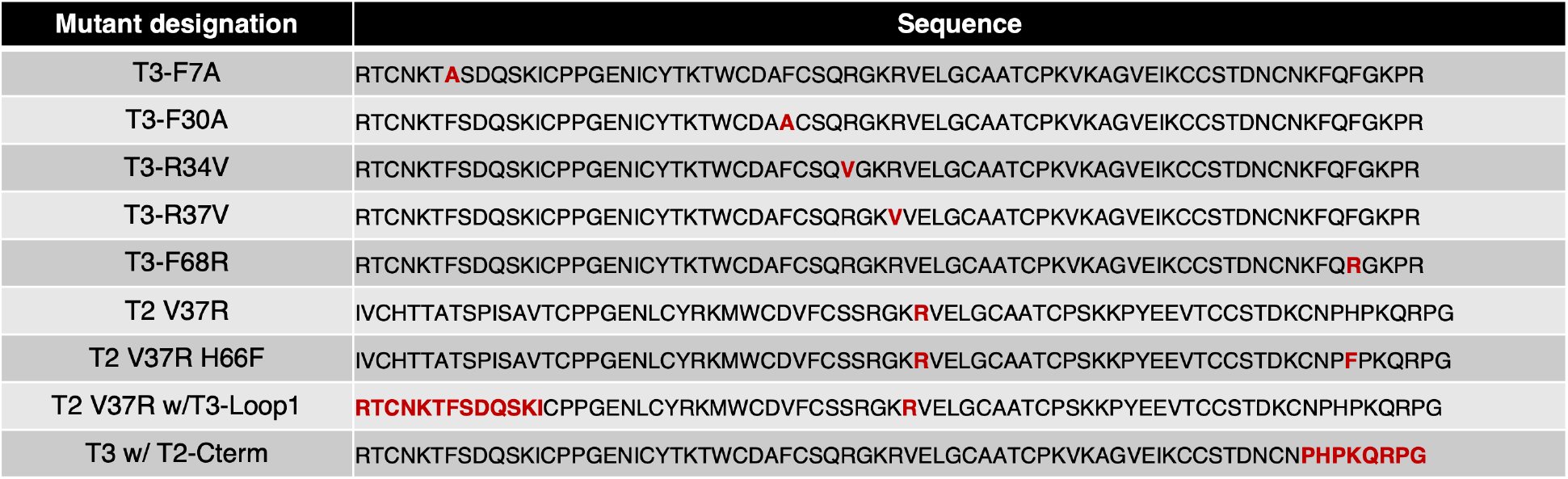
Mutated Toxin Sequences.

## Supplemental Methods

### Sculptor Algorithm

We define the following parameters:

*S*: Candidate set of interacting residues on the generated binder.
*R*: Full-atom coordinates of a target structure.
*F*: Interaction field.
*T_m_*: Metropolis Starting Temperature.
*γ*: Temperature Annealing Factor.
*α_z_*: Latent vector learning rate.
*α_ϕ_*: Transformation parameter learning rate.
*r*_in_, *r*_out_: Inner/Outer radius of initial position.
*∊*_refıne_: Fit cutoff for inclusion in the final sculpting loop.
*∊*_fınal_: Fit cutoff for including field residue identities for design.
*n*_interf_: Number of interface residues to optimize for.
*m*_field_: Number of outer-loop iterations to reassign the interface residues.
*m*_sculpt_: Number of inner-loop iterations to fit the target field residues.
*c*: Clipping value for SculptLoss (Described Below).

We use the following notation for variables:

*z*: Latent vector.
*ϕ*: 3D homogenous transformation parameters.
*x*: Cartesian coordinates.
*s*: Subset of binder residue positions in *C*.
*t*: Subset of interacting residues in *F*.
*ℓ*: Loss.

We define the following core functions:

*G*(*z*): Generative model with coordinate outputs.
*T*(*ϕ*): 3D homogenous transformation operation.

The pseudo-code of the Sculptor optimization loop of Sculptor is provided below:

#### Algorithm: Sculptor Optimization Loop.

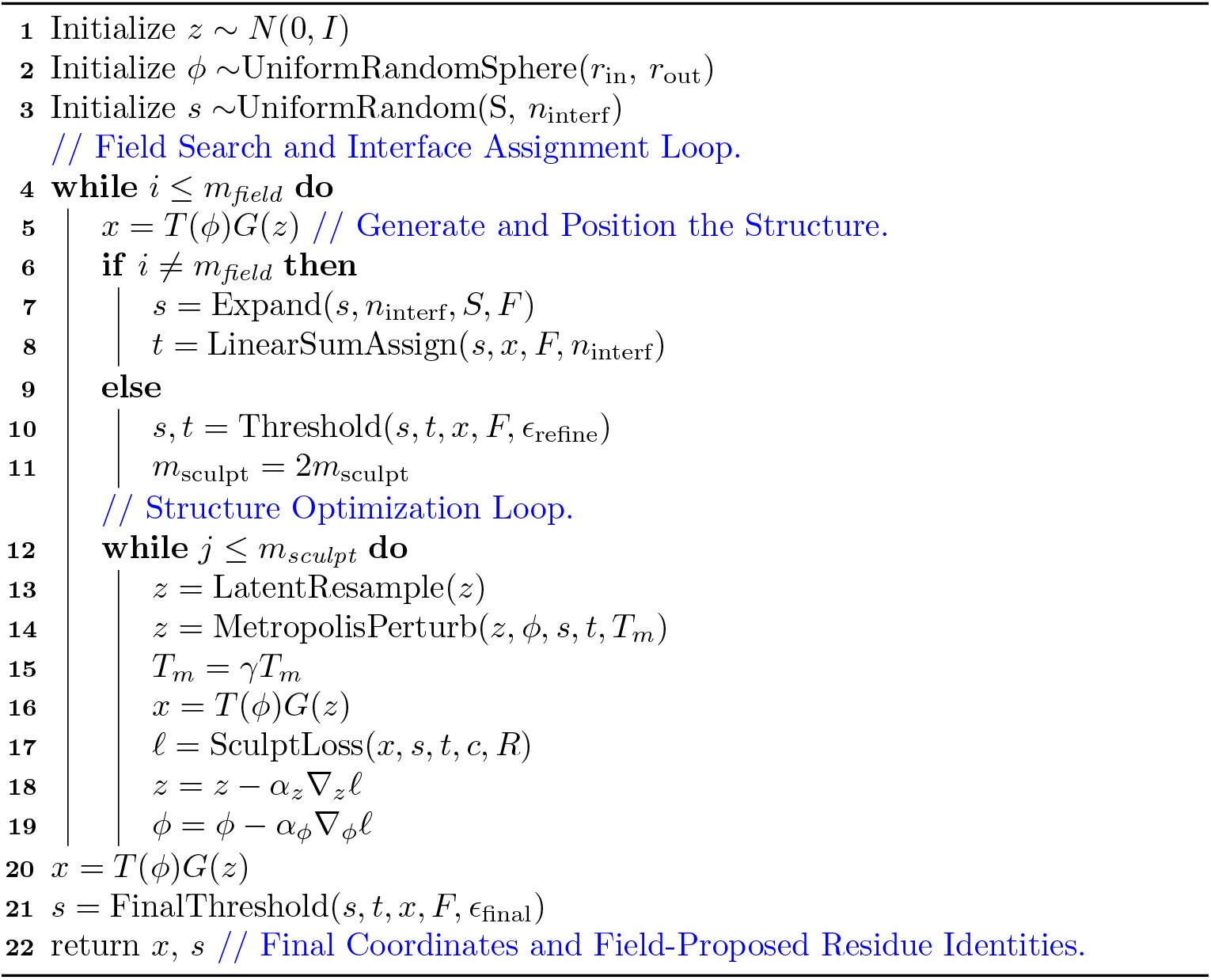

Descriptions of the functions used are provided below:

The *Expand* function is a stochastic sampler used to expand the number of residues in s to some number ≥ *n*_interf_ and ≤ |*S*|. In this study we expand to a size of 2*n*_interf_. The sampling distribution can be uniform random, or weighted based on the pre-computed ddG of a field interaction if available.

The *LinearSumAssign* function is used to assign pairs of interactions in *s* to the set of best fitting partners *t* ⊂ *F* without overlap. The loss used is the Frobenius norm between the *C_α_C_β_* vectors of the residues in *x* (specified by *s*), against those of the field residues. The function returns an updated *s*, and *t*, which are the target field residues for each backbone residue in *s*. The outputs *s* and *t* are both ordered, and contain *n*_interf_ elements. During the inner optimization loop, the algorithm tries to fit the residues specified by *s* into their target positions *t*. The linear sum assignment implementation can be found in the SciPy python package[1].

The *Threshold* function is used to remove any interacting pairs, specified by *s* and *t*, that fit the field worse than the threshold ∊_refıne_. The function returns the subset of *s* and *t* that satisfy this criterion. By doing this, the final structure optimization loop is used to improve the fit of well-fit residues, while discarding poor-fitting interactions.

The *LatentResample* function stochastically resamples elements of the latent vector based their distance from zero, to ensure that the optimization is restricted to parts of the latent distribution that are supported. This resampling scheme is based on prior work on latent vector recovery [2, 3], and the probability of resampling is given by the following function:

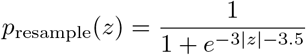

The *MetropolisPerturb* function stochastically perturbs the latent vector *z* by adding some vector ~ *N*(0, ∊*I*), and accepts or rejects the perturbation based on an increase or decrease in *SculptLoss* (described below)[4, 5]. The acceptance probability is determined by the following criterion:

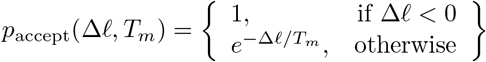

The *SculptLoss* function is described in the next section.

### Sculptor Loss Function

*SculptLoss* is the primary loss function in the optimization loop, and consists of a repulsive term and a fitting term that are both fully differentiable and empirically tuned for stability. The two terms are summed to compute the full loss function:

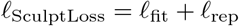

### Repulsive Loss

Let *d*_vdw_ denote the sum of the van der Waals radii of an atom pair, *σ* the sigmoid function, and ReLU the Rectified Linear Unit. The repulsive term is given by:

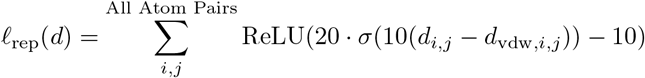

For the sculpted molecule, each residue is treated as an alanine, while the target is treated as full-atom, including side-chain.

### Fitting Loss

The fitting loss term is a *L*_2_ loss between the generated and target *C_α_C_β_* vectors, where each element is weighted by the inverse of the *L*_2_ loss for that residue pair, and clipped via the parameter *c*. Let the *C_α_C_β_* coordinate vectors of the *n*_interf_ residues in *s* and *t* be denoted as *v_s_* and *v_t_*. We compute the fitting loss as follows. First, we compute the squared *L*_2_ loss vector *l* with each element is given by:

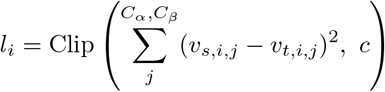

The *l* vector is then used to compute a normalized weight vector *w* with the following elements:

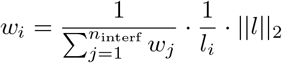

The weight vector is then used to re-weight the loss vector, which is summed over all residues in the interface set:

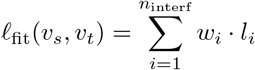

Notably, the fitting loss and its weighting plays a crucial role in Sculptor, adding an element of “stickiness” to the algorithm behavior. Specifically, residues that fit field interactions well are retained and further optimized, while poorly fitting ones are allowed to drift and are more likely to be reassigned. This tendency significantly improves the quality of field fitting, as well as interface search efficiency.

